# A CheZ orthologue in *Campylobacter jejuni* plays a role in chemotaxis through conserved phosphatase activity

**DOI:** 10.1101/2023.01.06.523011

**Authors:** Abdullahi S. Jama, Julian M. Ketley

**Author notes:** **Corresponding author** Prof Julian Ketley, Department of Genetics and Genome Biology, University of Leicester, University Road, Leicester, LE1 7RH UK.

## Abstract

The major food-borne pathogen *Campylobacter jejuni* employs chemotactic motility to colonise the avian gut, and also as a virulence mechanism in human diarrhoeal disease.

In *Escherichia coli* CheY activity is modulated by CheZ, a phosphatase originally thought to be absent in *C. jejuni*. The Hp0170 protein of *Helicobacter pylori* is a distant homologue of CheZ and, as *C. jejuni* Cj0700 is homologous to HP0170, Cj0700 could also act as a CheZ orthologue in *Campylobacter*. Both the *C. jejuni* CheV and CheA proteins also contain a response regulator (RR) domain that may be phosphorylated. Cj0700 would therefore be predicted to dephosphorylate *C. jejuni* CheY and possibly also the CheV and CheA RR domains.

A mutant (Δ*cj0700)* and complement (Δ*cj0700, cj0046*::*cj0700*) were constructed in *C*.*jejuni* strains NCTC11168, NCTC11828 and 81-176. On semisolid agar the Δ*cj0700* mutant strain showed reduced motility relative to wild-type and this phenotype was reversed in the complemented strain. In pull down and bacterial two hybrid assays, expressed Cj0700 was able to interact with CheY, CheA-RR and CheV.

Cj0700 is able to dephosphorylate the RR domain of CheY and CheA-RR, but less efficiently, CheV. These findings verify that Cj0700 plays a role in *C. jejuni* chemotaxis through phosphatase activity with respect to CheY, and is hence likely to be a CheZ orthologue. Cj0700 also partially modulates the phosphorylation level of the RR domain on CheA and CheV, although the functional consequences of this interaction require further investigation.

## Introduction

*Campylobacter jejuni* is a spirally-curved zoonotic pathogen of significant clinical importance (1) that has been identified as a high priority pathogen by WHO (2). *C. jejuni* is responsible for high levels of morbidity through food-borne gastroenteritis, with rising numbers in developed nations and endemic status worldwide (3). It is also associated with complications such as Guillain-Barré syndrome (4). Despite an enormous burden on world health, understanding of the pathophysiological mechanisms of disease remain ill-defined. The main reservoir for transmission to humans is the intestine of warm-blooded animals and birds, which are not always colonised asymptomatically (5). Reduction of intestinal colonisation, particularly avian, is one approach to reduce the impact of campylobacters on human health.

The *C. jejuni* cell has a polar flagellum at one or both ends which, combined with the spirally curved morphology, provides a high degree of screw-like darting motility. Chemotaxis is a key colonisation factor, enabling migration towards favourable environments and evasion of toxic agents (6). This behaviour is implicated in virulence, involving a direct relationship between chemotactic motility and chemical components directly involved in infection (7-10). A detailed understanding of the sensory and transduction mechanisms of chemotactic motility in *C. jejuni* is essential for the understanding of this pathogen’s ability to cause disease.

Chemotactic motility involves a series of signal transduction proteins that transduce the status of the extracellular chemical environment to the flagellar motor resulting in differential rotation (11). The backbone of this transduction system consists of the CheA, CheW and CheY proteins. CheA and CheW form part of chemoreceptor complexes at the cell surface (12). Changes in ligand-chemoreceptor interaction state results in autophosphorylation of CheA, a histidine protein kinase (HK). CheA transfers the phosphate group to the response regulator (RR) domain of CheY, which then interacts with the flagellar motor to modulate flagellar rotation (13); in *Escherichia coli* this occurs in the absence of ligand-receptor binding. Rapid modulation of flagellar rotation in response to changing environments involves receptor adaptation via methylation/demethylation by CheR and CheB (14, 15) and the rapid removal of phosphate from CheY. CheY has an inherent dephosphorylation activity, but other mechanisms including phosphatases, for example CheZ (16), and transfer of phosphate to multiple CheY homologues (a phosphate sink), have been linked to phosphate removal from CheY (11).

The *C. jejuni* flagellar system is an essential virulence determinant both through its role in motility (9, 17) and also through the flagellar Type Three Secretion System (T3SS) that has a role in the secretion of invasion proteins (18). In addition to transducer-like protein chemoreceptor variation (6, 7, 19), the chemotaxis signal transduction pathway of *C. jejuni* is unique, with a combination of differences linked to mechanisms that likely modulate the activity of pathway components. CheA, similar to some other bacteria, has a RR domain (CheA-RR) in addition to the HK domain (CheA-HK), there is a CheV component that consists of a RR domain (CheV-RR) linked to a CheW-like domain, and CheB lacks a RR domain. Like *Helicobacter pylori, C. jejuni* expresses ChePep, a novel chemotaxis regulator unique to *Epsilonproteobacteria* (20), but was thought to lack a functional homologue of CheZ (21, 22).

As *C. jejuni* CheV, which is involved in coupling CheA to chemoreceptors, and CheA have RR domains, it is possible that, like CheY, these domains are phosphorylated by the CheA-HK domain. These additional RR domains could act as a phosphate sink to modulate CheY phosphorylation as an alternative to CheY dephosphorylation by CheZ, but studies in the closely related *H. pylori* highlighted the presence of a previously uncharacterised (23) remote homologue of CheZ (HP0170) (24) that has phosphatase activity similar to CheZ (25). This homologue is also present in *C. jejuni* NCTC11168 (*cj0700*; 49.4% identity) (24).

Here we show that in addition to CheY, *C. jejuni* CheA is able to phosphorylate both the CheA-RR and CheV-RR domains. *C. jejuni* also encodes a *cheZ* orthologue (*cj0700*) which we show dephosphorylates CheY and plays a role in the chemotaxis phenotype of *C. jejuni*.

## Materials and methods

### Bacterial strains and media

*C*.*jejuni* strains (Table 1) were cultured microaerobically at 42°C using Mueller-Hinton agar (MHA) or broth (MHB) supplemented with vancomycin (10 μg/ml) and trimethoprim (5 μg/ml). *E. coli* (Table 1S) was grown on Luria-Bertani or MacConkey media at 37°C. Where necessary, selection using media supplemented with appropriate antibiotics was used (supplementary methods).

**Table 1:** *Campylobacter* strains used in this study.

| Name | Description | Source/Reference |
| --- | --- | --- |
| NCTC 11168 | Wild type | NCTC, Colindale, London, UK |
| NCTC 11828 | Wild type also known as 81116 | (Manning et al., 2001) |
| 81-176 | Wild type | (Korlath et al., 1985) |
| AJ1 | NCTC 11168 $\Delta cj0700::cat$ (Cm<sup>R</sup>) | This study |
| AJ4 | NCTC 11828 $\Delta cj0700::cat$ (Cm<sup>R</sup>) | This study |
| AJ6 | 81-176 $\Delta cj0700::cat$ (Cm<sup>R</sup>) | This study |
| AJ3 | NCTC 11168 $\Delta cj0700::cat$, $cj0046::cj0700$ (Km<sup>R</sup> & Cm<sup>R</sup>). | This study |
| AJ7 | NCTC11828 $\Delta c8j\_0667::cat$, $cj0046^*::cj0700$ (Km<sup>R</sup> & Cm<sup>R</sup>). | This study |
| AJ10 | 81-176 $\Delta cjj81176\_0723::cat$, $cj0046^*::cj0700$ (Km<sup>R</sup> & Cm<sup>R</sup>). | This study |
| 11168cheY | *cheY* mutant of NCTC11168 (Km<sup>R</sup>) | (Elgamoudi et al., 2018) |
| *cj0046\** is used as a label for the homologous region in NCTC11828 and 81-176. |  |  |

### Chemotaxis and Growth Assays

Basic cell motility of *C. jejuni* strains was determined by phase contrast microscopy and chemotaxis by inoculating *C*.*jejuni* cell suspensions onto semi-solid MHA plates (26). *C. jejuni* growth patterns were determined either in individual tubes or in microtiter plates in a FLUOstar Omega microplate reader. Both involved incubation at 42°C with shaking under microaerobic conditions (supplementary methods).

### *C. jejuni* mutant construction

The *cj0700* gene, or its homologue, of *C. jejuni* strains NCTC11168, NCTC11828 and 81-176 (Table 1) were mutated by insertion of a promoter-less and terminator-less chloramphenicol acetyltransferase (*cat*) gene, and deletion of approximately 700 bp of the *cj0700* coding sequence (Table 1 and supplementary methods). For complementation of the *cj0700* mutation, the wild-type *cj0700* allele, under the control of the *C. jejuni metK* promoter, was inserted into the pseudogene *cj0046* in NCTC11168 and the equivalent locus in other strains (Table 1). Mutant loci were transferred into motile variants of each strain by natural transformation and motility verified (supplementary methods). Mutations were confirmed by PCR and DNA sequencing.

### Protein expression and purification

His-tagged CheY, CheA, CheA-HK, CheA-RR CheV and Cj0700, and GST-tagged Cj0700 and CheY were expressed from *E. coli* using IPTG induction, followed by column purification, as appropriate (supplementary methods). CheA-HK and CheA-RR consisted of the isolated kinase and response regulator domains from CheA, respectively.

### Phosphorylation/de-phosphorylation assays

Phosphorylation/de-phosphorylation assays were carried out using γ^32^-P ATP diluted with unlabelled ATP. CheA-HK was incubated with radio-labelled Pi prior to addition of RR domain-carrying or Cj0700 proteins. For protein-protein interaction assays of phosphorylated protein, GST-tagged protein was bound to glutathione Sepharose 4B resin beads and His-tagged protein added; protein was phosphorylated by the addition of acetyl phosphate. See supplementary methods for details.

### Bacterial two hybrid assays

The bacterial two hybrid (B2H) assay was based on adenylate cyclase (AC) reconstitution (27) (supplementary methods). Interactions were screened on MacConkey agar containing 1% (w/v) glucose-free maltose incubated at 30 °C and colour change was evaluated visually.

## Results

### Mutation of cj0700 affects chemotactic motility

The potential *cj0700* phosphatase gene in NCTC11168 and the predicted homologues in NCTC11828 (C8J_0667) and 81-176 (CJJ81176_0723) were mutated (Table 1; AJ1, AJ4 and AJ6, respectively). All three *cj0700* mutants were complemented by insertion of *cj0700* into the pseudogene *cj0046*, or its equivalent region in NCTC11828 and 81-176 (Table 1; AJ3, AJ7 and AJ10, respectively). The pattern of chemotactic motility of the different mutant strains was then determined.

The swarm assay reflects chemotactic motility as the cells move away from the point of inoculation along a nutrient concentration gradient. A NCTC11168 *cheY* mutant (11168cheY, Table 1) displayed less chemotactic motility when compared to the wild-type, which showed a characteristic ring pattern of spread (Figure 1A). Mutation of *cj0700* causes a defect in chemotactic motility when compared to wild type (Figure 1BCD). Loss of chemotaxis was observed in all three strain backgrounds, and this was restored in the complemented strains with some variation (Figure 1BCD).**D**

**Figure 1.**
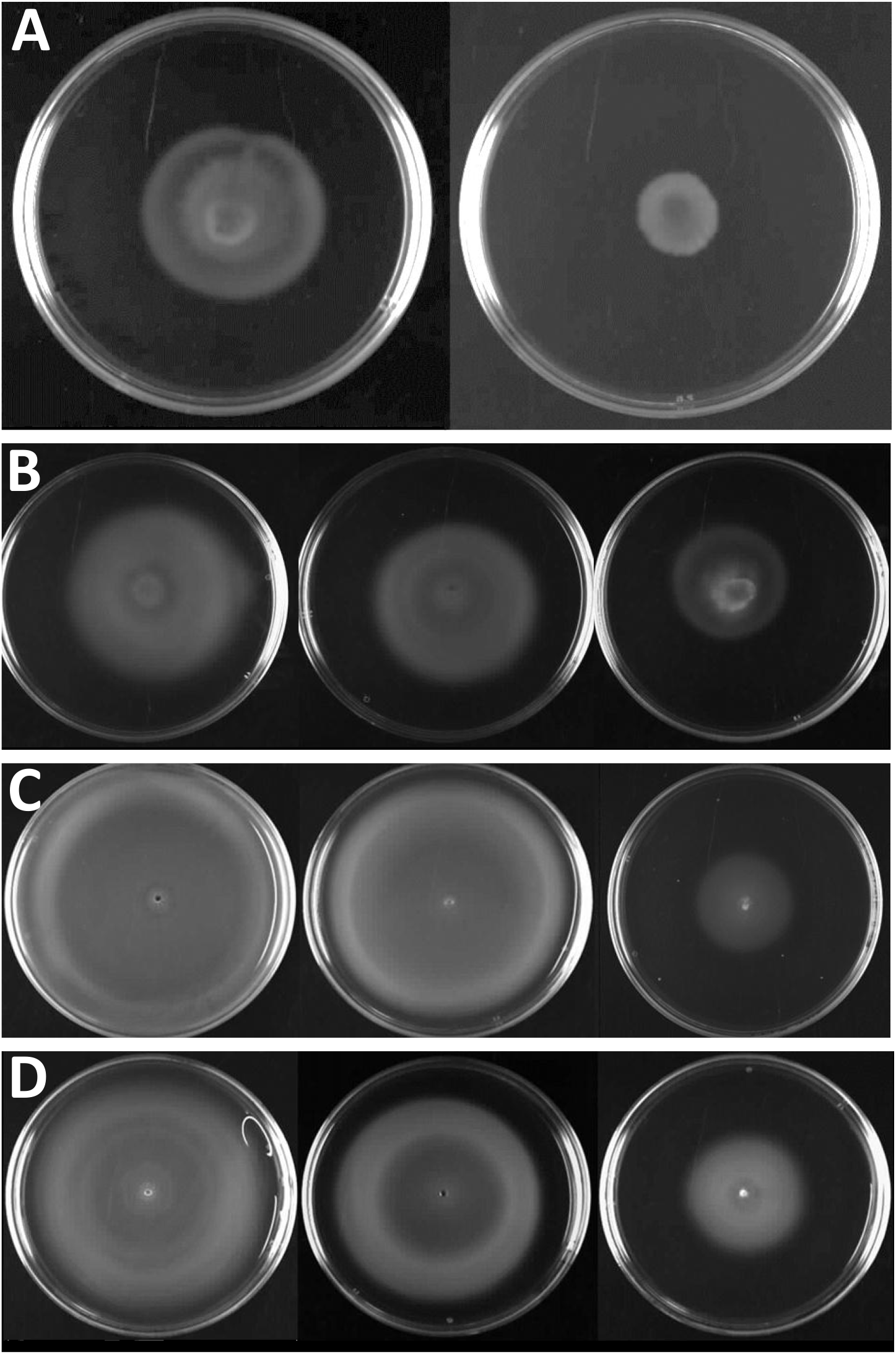
Chemotactic motility of *cj0700* mutants and their complements in different strain backgrounds. Cells were inoculated into the centre of 0.3% (w/v) MHA and grown for 72 hours at 42 °C in a microaerobic atmosphere. (A) NCTC11168 and 11168cheY; (B) NCTC11168, AJ3 and AJ1; (C) NCTC11828, AJ7 and AJ4; (D) 81-176, AJ10 and AJ6.

### Cj0700 interacts with CheY, CheA-RR and CheV

To determine if Cj0700 can bind to CheY, CheA-RR or CheV proteins, protein-protein interactions were investigated using pull down experiments and then confirmed by a bacterial two hybrid system (B2H).

We utilised an *in vitro* GST-based, protein-protein interaction pull-down using GST and His-tagged proteins to show that Cj0700 can interact with phosphorylated CheY and CheA-RR (Figure 2). His-tagged Cj0700 binds to GST-tagged CheY, phosphorylated with acetyl-phosphate and immobilised onto glutathione beads (Figure 2, lane 3), but not with GST alone (Figure 2, lane 4). His-tagged CheA-RR also interacts with immobilised CheY, although some nonspecific interaction with GST and both His-CheA (Figure 2, lane 2) and His-CheV (data not shown) was observed. Given these non-specific interactions, an alternative approach of a two-hybrid system was used to investigate the ability of Cj0700 to interact with CheA-RR and CheV and verify that seen between Cj0700 and CheY using pull-down experiments (Figure 2).

**Figure 2.**
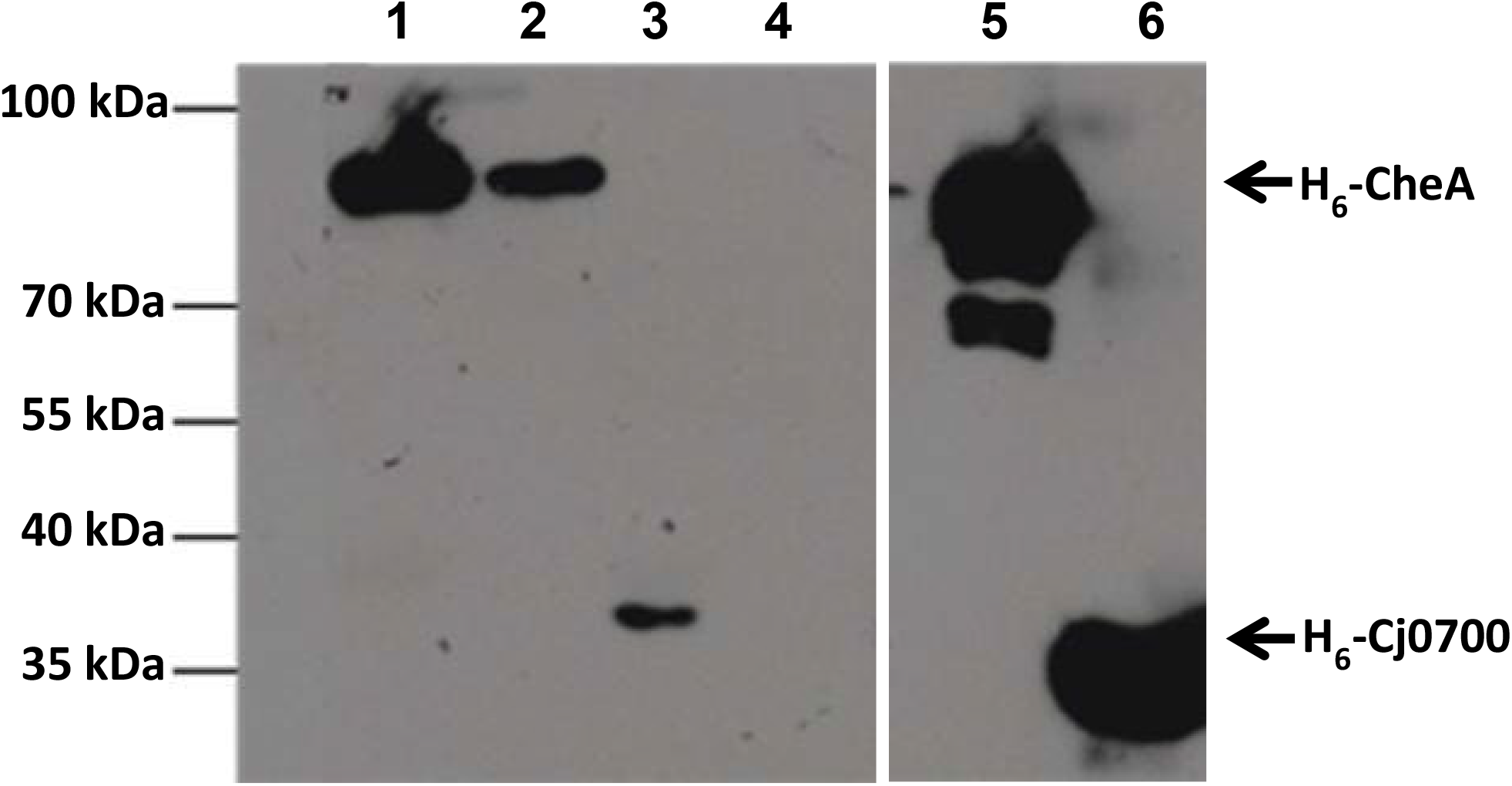
Determination of protein-protein interactions between Cj0700 with Che proteins. In vitro protein-protein interaction pull-down experiment using glutathione beads and GST- and His-tagged Cj0700 and Che proteins. Western blot analysis of His-tagged Cj0700 protein pull-down with GST-tagged CheY protein in the presence of Mg^2+^ and acetyl-phosphate. Detection of Cj0700 interaction with CheY by anti-His HRP conjugate antibody. GST-CheY+His-CheA (lane 1), His-CheA+GST (lane 2), His-Cj0700 + GST-CheY (lane 3), His-Cj0700 + GST (lane 4), His-CheA (lane 5), and His-Cj0700 (lane 6). Pull down reactions shown in lanes 5 and 6 did not contain Mg^2+^ and in the reactions shown in lanes 1 and 3 GST-CheY was phosphorylated with acetyl phosphate.

To use an adenylate cyclase (AC)-based two hybrid system (27), Cj0700 was fused to the T18 AC subunit and CheY, CheA-RR, and CheV were each fused to the T25 subunit (Table 2). A leucine zipper-based interaction (pT25:Zip/p18:Zip) (27), used to confirm interaction of two fragments of AC, was detected using MacConkey agar. The ability of the CheA-RR fragment to dimerise showed an expected strong positive interaction while the lack of a RR domain in CheB was reflected in the absence of colour change on MacConkey agar. There was a strong interaction observed between Cj0700 and CheY, but no interaction with CheB. The interaction of Cj0700 with CheA-RR or CheV in the B2H system was found to be moderate compared to CheY or the positive control. These data confirm that Cj0700 interacts with CheY, but the interaction with CheA-RR or CheV appears less strong.

**Table 2.** Bacterial Two Hybrid screen of Che proteins and Cj0700.

| T18 AC subunit | T25 AC subunit | Interaction strength |
| --- | --- | --- |
| T18:Zip | T25:Zip | ++ |
| T18:A-RR | T25:A-RR | ++ |
| T18:Cj0700 | T25:CheB | - |
| T18:A-RR | T25:Zip | - |
| T18:Cj0700 | T25:CheY | ++ |
| T18:Cj0700 | T25:CheV | + |
| T18:Cj0700 | T25:CheA-RR | + |
| Interaction strength reflected colour change on MacConkey media. ++ strong interaction, + weak interaction and – no interaction. |  |  |

### C. jejuni CheA mediates phosphotransfer to CheY, CheA-RR domain and CheV

As a prelude to characterizing the phosphate transfer between CheA and CheA-RR or CheV-RR, the fundamental ability of *C. jejuni* CheA to autophosphorylate and subsequently mediate phosphotransfer to CheY was established. GST- and His-tagged purified proteins were used in the kinase assays. In the presence of γ^32^-P ATP CheA is able to stably autophosphorylate (data not shown), however in the presence of CheY neither protein was observed to be phosphorylated (data not shown). As native CheA containing the CheA-RR domain was unsuitable for phospho-transfer experiments under the assay conditions, the isolated CheA-HK domain was used. Autophosphorylated (γ^32^-P) CheA-HK was able to transfer phosphate to CheY (Figure 3), whereas in the absence of CheY, CheA-HK remained stably phosphorylated for the duration of the assay. Therefore, as predicted, *C. jejuni* CheA can phosphorylate CheY.

**Figure 3.**
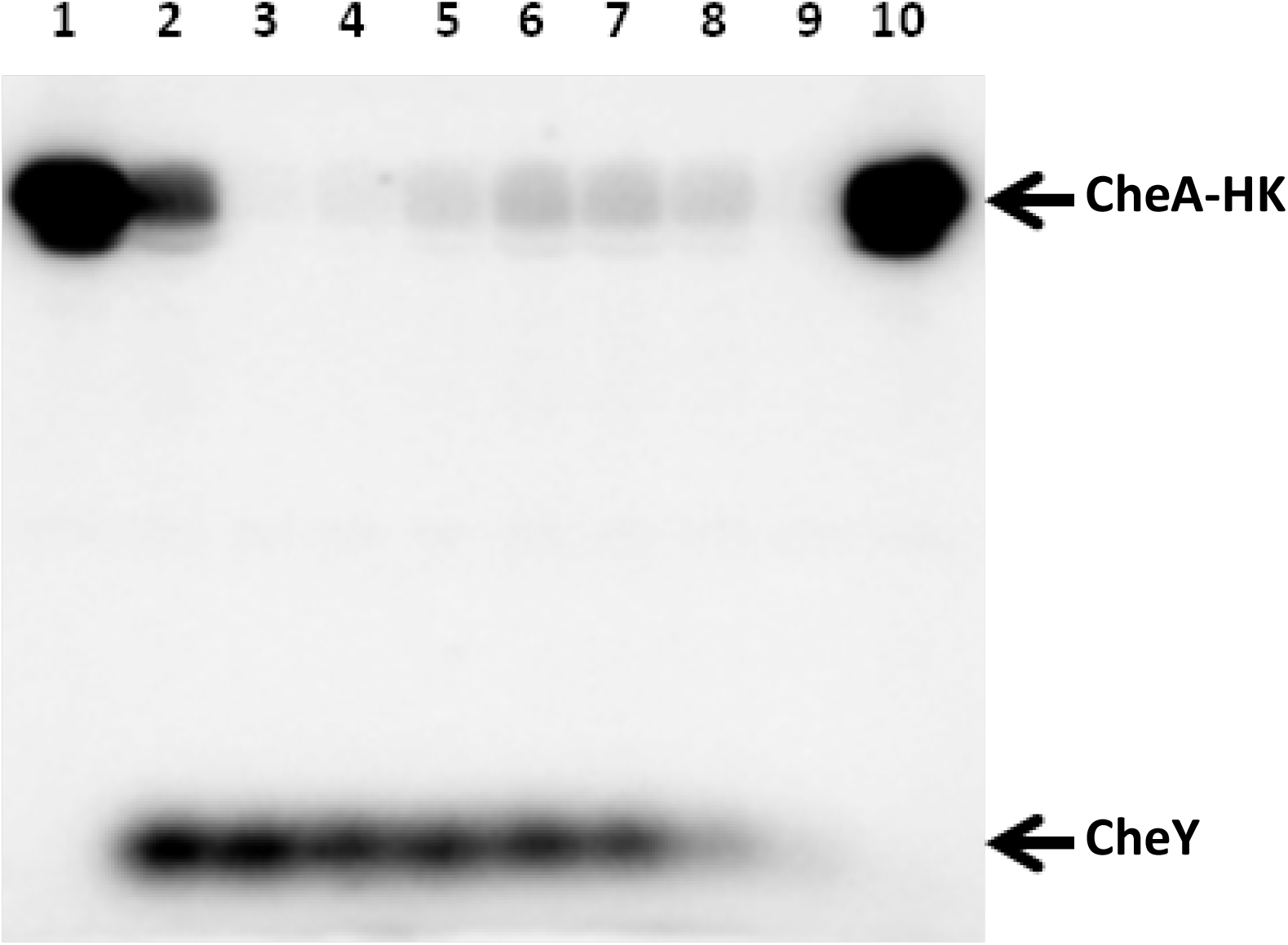
CheA-HK mediated transfer of phosphate to CheY. Phosphor screen visualisation of kinase assay between 40 μM CheA-HK and 15 μM CheY at 4°C. CheA-HK was incubated with 3 μM [γ-^32^P] ATP diluted with 30 μM non-radiolabelled ATP for 10 minutes before reduction of the reaction temperature to 4°C. A sample was taken as an initial CheA-HK autophosphorylation control (Lane 1) and another sample was incubated separately under assay conditions for 10 minutes (Lane 10). CheY was added to the remaining reaction master mix and a sample taken immediately (Lane 2), then at 20 and 40 secs, 1, 3, 4, 5 and 10 mins (Lanes 3-9).

*C. jejuni* CheA contains a RR domain and although it is likely that it is able to be phosphorylated by CheA-HK, this has not been demonstrated. Autophosphorylated (γ^32^-P) CheA-HK was observed to phosphorylate CheA-RR immediately, with the level of phosphorylation then declining steadily (Figure 4A). Complete Pi hydrolysis was not observed after 30 minutes, as a low level of labelled CheA-RR was still observable. As before, in the absence of CheA-RR, CheA-HK retained label, indicating that loss of phosphate from CheA-HK in the presence of CheA-RR was due to transfer of phosphate to CheA-RR and not solely because of CheA-HK autodephosphorylation. CheV has a RR domain which may modulate CheV function via phosphorylation state. CheA is one kinase that may have a phospho-transfer role to CheV. Autophosphorylated (γ^32^-P) CheA-HK was able to phosphorylate CheV, but CheV dephosphorylated by 30 seconds (Figure 4B). CheA-HK autophosphorylation and subsequent loss of phosphate in the presence of CheV indicates that CheA-HK is able to transfer ^32^P to the RR domain of CheV.

**Figure 4.**
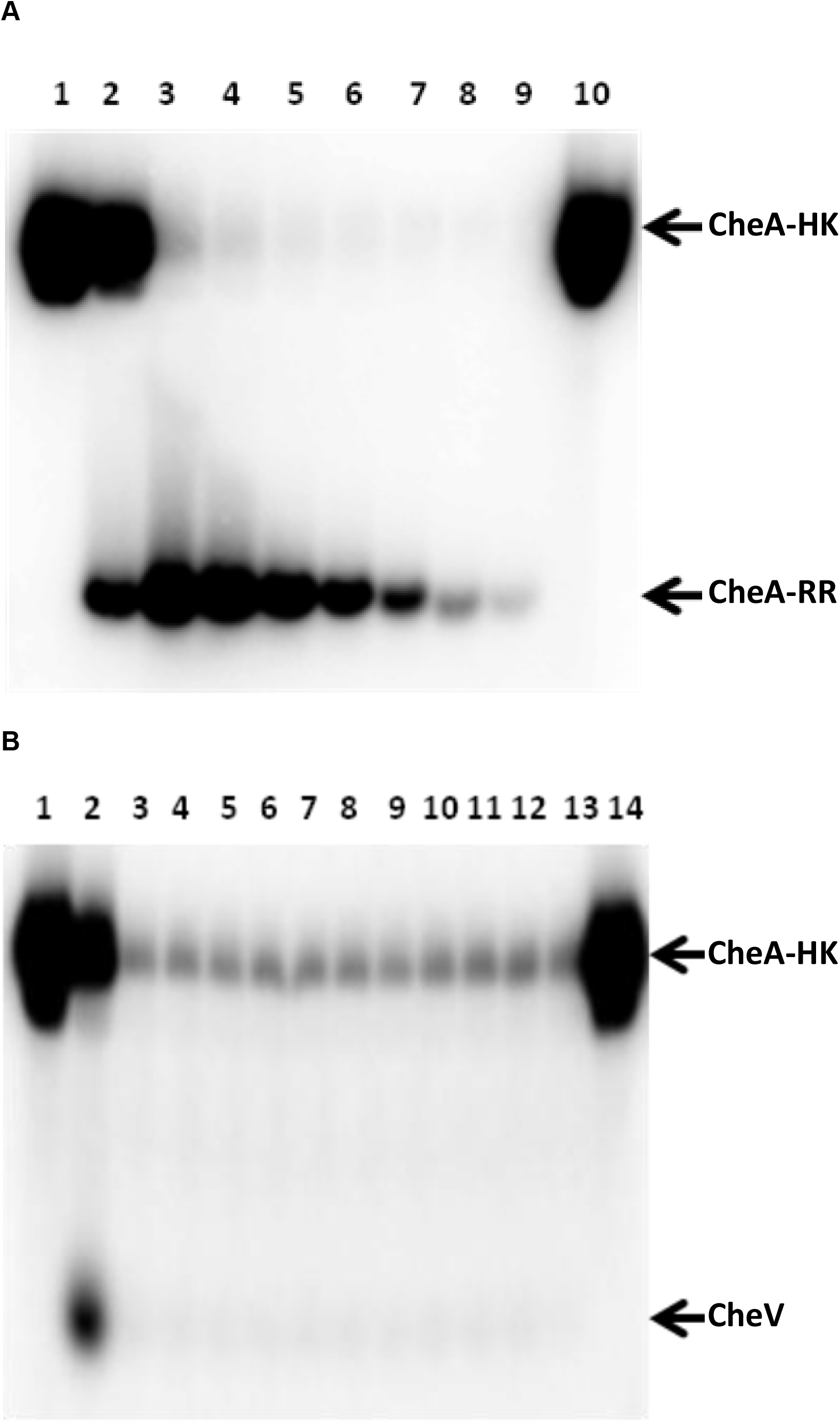
A & B. CheA-HK mediated transfer of phosphate to CheA-RR and CheV. Phosphor screen visualisation of kinase assay between 20 μM CheA-HK and 20 μM CheA-RR (A) and 20 μM CheA-HK and 20 μM CheV (B). CheA-HK was incubated with 1.3 μM [γ- ^32^P] ATP diluted with 30 μM non-radiolabelled ATP for 10 minutes before incubation at the reaction temperature of 42°C. A sample was taken as an initial CheA-HK autophosphorylation control (Lane 1) and another sample was incubated under assay conditions for the duration of the experiment (A: Lane 10 and B: Lane 14). CheA-RR (panel A) was added to the remaining reaction master mix and a sample was taken immediately (Lane 2) then at 2, 5, 10, 15, 20, 25 and 30 minutes (Lanes 3-9). CheV (panel B) was added to the remaining reaction master mix and a sample was taken immediately (Lane 2) then at 0.5, 1, 1.5, 2, 2.5, 3, 3.5, 4, 4.5, 5, and 10 minutes (Lanes 3-13).

### Cj0700 de-phosphorylates response regulator domains of CheA and CheY, but not CheV

CheA-HK can phosphorylate CheY, CheA-RR, and CheV in *C. jejuni*. Phosphorylation assays also show that each RR domain showed differing levels of auto-dephosphorylation under assay conditions. In order to determine if Cj0700 is a functional homologue of CheZ, the phosphatase activity of Cj0700 towards the Che domains phosphorylated by CheA was investigated. Cj0700 was expressed and the dephosphorylation of CheY phosphorylated by CheA-HK was determined (Figure 5). Initially, CheA-HK was incubated in reaction buffer with γ^32^-P-ATP. Again, CheA-HK, autophosphorylated with γ-^32^P subsequently transferred phosphate to CheY with a peak intensity at 90 seconds (Figure 5). In the presence of Cj0700, removal of phosphate from CheY occurred at a greater rate than phosphate was donated by CheA-HK. It is unlikely that Cj0700 blocks transfer of phosphate from CheA-HK to CheY, as both in the presence and absence of Cj0700 there is an equivalent loss of signal from auto-phosphorylated CheA-HK at all time points.

**Figure 5.**
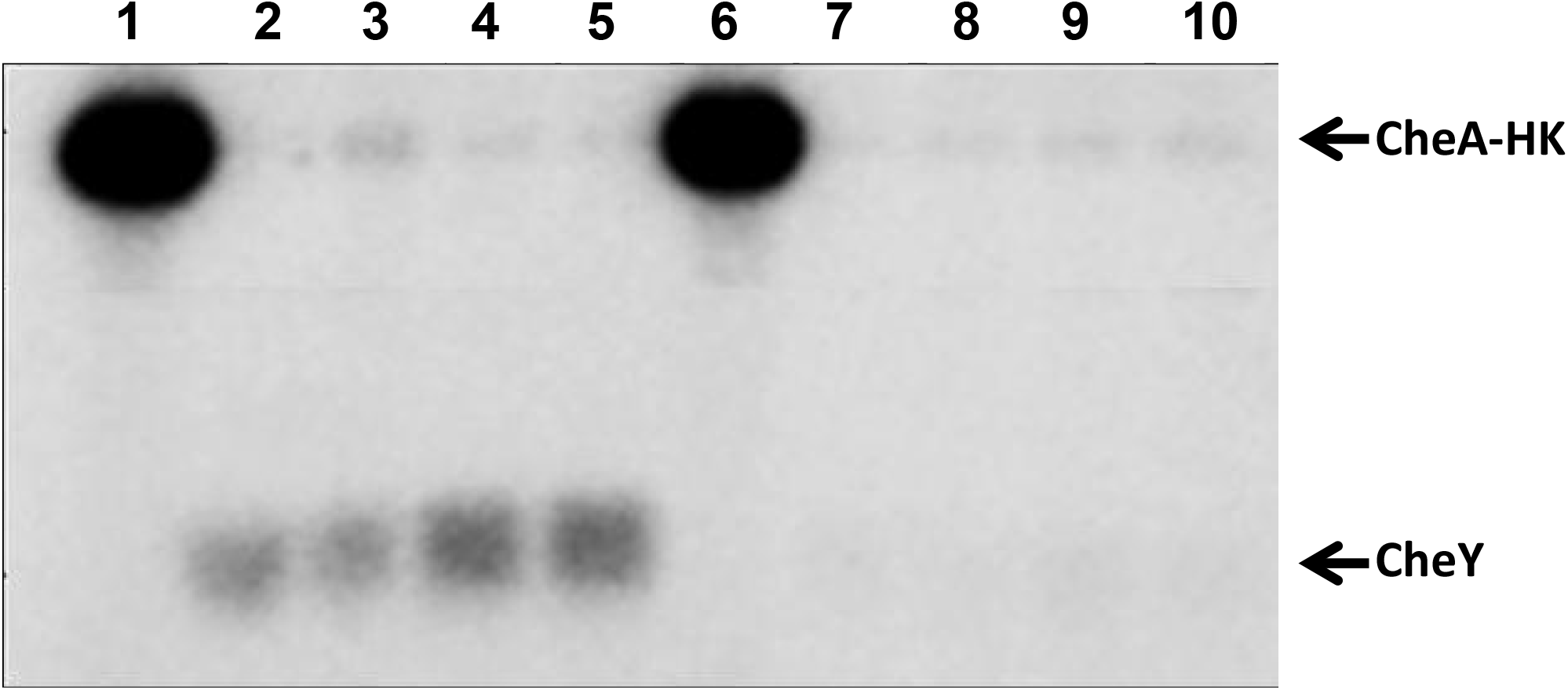
Cj0700 releases phosphate from phosphorylated CheY. Phosphor screen visualisation of kinase assay between 5 μM CheA-HK and 15 μM CheY at 30°C. CheA-HK was incubated with 3 μCi [γ-^32^P] ATP and 5 mM non-radiolabelled ATP for 10 minutes. A sample was taken as an initial CheA-HK autophosphorylation control (Lanes 1 and 6). CheY was added to the remaining reaction master mix without (lanes 2-5) and with 15 μM Cj0700 (lanes 7-10). Samples were taken immediately (Lanes 2 and 7) then at 30, 60, and 90 seconds (Lanes 3-5 and 8-10).

As Cj0700 can dephosphorylate CheY, Cj0700-mediated de-phosphorylation of CheA-RR was investigated. Phosphate (γ-^32^P) transferred from CheA-HK to CheA-RR is removed in the presence of Cj0700 over a 30-minute time course (Figure 6A). Although Cj0700 does de-phosphorylate CheA-RR, this occurs markedly more slowly than with phosphorylated CheY (Figure 5). In a similar experiment, the ability of Cj0700 to de-phosphorylate CheV following phosphorylation by CheA-HK was determined. Phosphate (γ-^32^P) transferred by CheA-HK to CheV does not appear to be removed in the presence of Cj0700 over a 90 second time course (Figure 6B). Therefore, in contrast to CheY and CheA-RR, Cj0700 did not detectably remove phosphate from CheV under the conditions tested.

**Figure 6.**
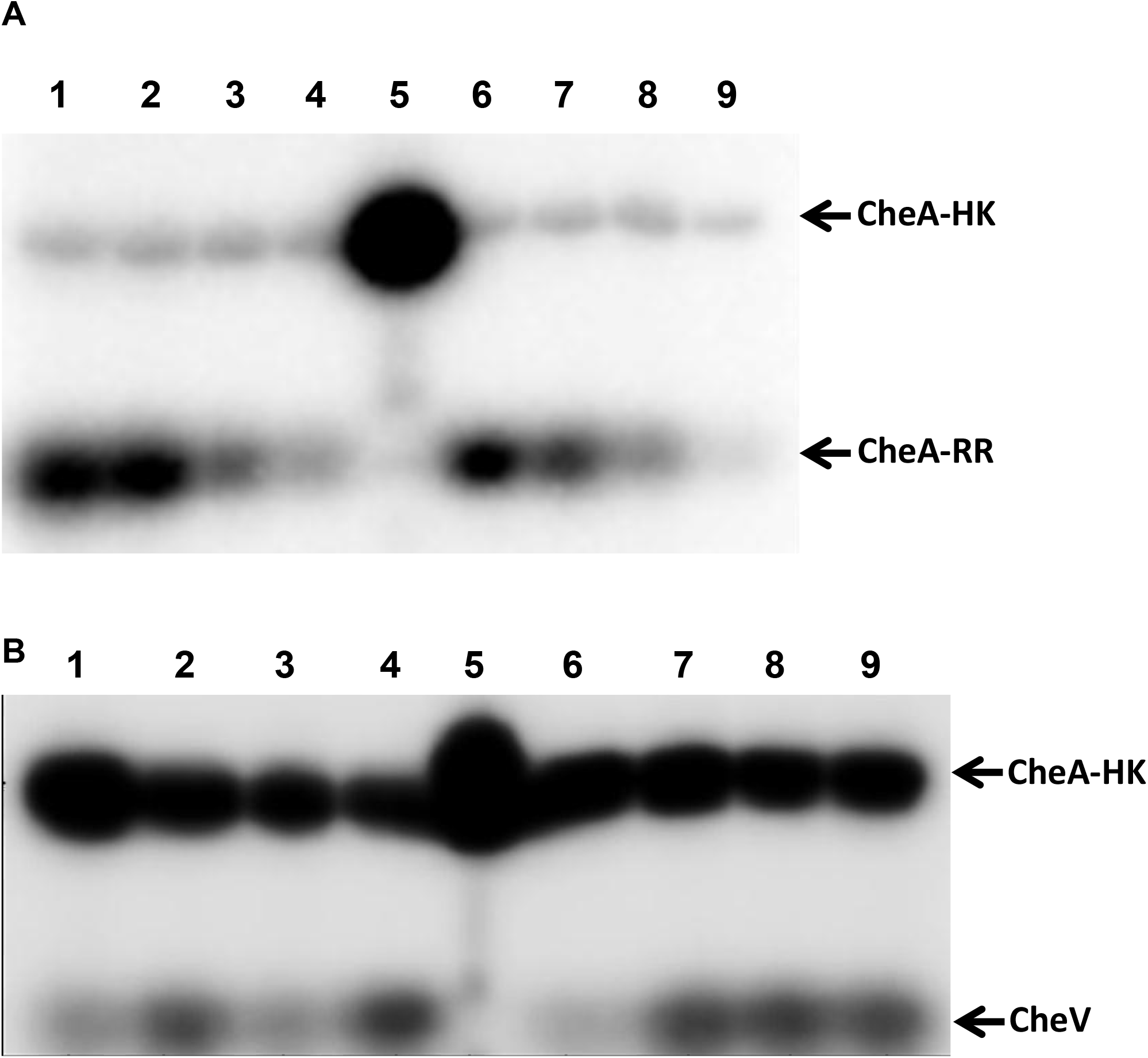
A & B. Cj0700 releases phosphate from phosphorylated CheA-RR but not from CheV. Phosphor screen visualisation of kinase assay between 5 μM CheA-HK and 5 μM CheA-RR at 30°C (A) and CheA-HK and 10 μM CheV at 4°C (B). **Panel A**. CheA-HK was incubated with 3 μCi [γ-^32^P] ATP and 5 mM non-radiolabelled ATP for 10 minutes. A sample was taken as an initial CheA-HK autophosphorylation control (Lane 5). CheA-RR was added to the remaining reaction master mix without (lanes 1-4) and with 15 μM Cj0700 (lanes 6-9). Samples were taken at 5, 10, 20 and 30 minutes (Lanes 1-4 and 6-9). **Panel B**. CheA-HK was incubated with 3 μCi [γ-^32^P] ATP and 5 mM non-radiolabelled ATP for 10 minutes at 30°C. A sample was taken as an initial CheA-HK autophosphorylation control (Lane 5). CheV was added to the remaining reaction master mix without (lanes 1-4) and with 15 μM Cj0700 (lanes 6-9). Samples were taken at 0, 30, 60 and 90 seconds (Lanes 1-4 and 6-9).

## Discussion

Chemotactic motility in *C. jejuni* has a pivotal role in intestinal colonisation (28) and the invasion of epithelial cells (29). The Che proteins responsible for signal transduction between the surface signal receptors and the flagellar motor complex in *C. jejuni* are notably different to the *E. coli* paradigm (30). These differences include the presence of a CheV protein, response regulator domain differences in CheB and CheA, and addition of modulating proteins that are absent in *E. coli* (ChePep) (20). Previously it was believed that *C. jejuni* also lacked a homologue of the phosphatase, CheZ, that in *E. coli* dephosphorylates CheY, affecting binding of CheY to the motor complex (23). CheZ was thought to be specific to members of the *beta* and *gamma proteobacteriacae*. In *H. pylori*, Hp0170 has been confirmed as a remote homologue of CheZ having been identified as a suppressor of non-chemotactic *cheW* mutants (24). The closest homologue of Hp0170 in *C. jejuni* is *cj0700* and therefore it was suggested that *C. jejuni* also controls the level of CheY phosphorylation through dephosphorylation using a CheZ-like homologue (25). Here we have shown that Cj0700 has the functional attributes expected of a CheZ homologue in that it is required for chemotactic motility and can bind to and dephosphorylate CheY.

A functional role in chemotactic motility was determined by testing the behaviour of a *cj0700* mutant in a well-established motility assay (26). Mutation of *cj0700* severely compromised the ability of *C. jejuni* to spread across semi-solid agar in response to a nutrient gradient established due to microbial growth. Given that the chemotaxis assay requires growth of the inoculated cells as they spread across the agar, we verified that the disruption of chemotaxis was not due to a growth defect resulting from the insertional mutation of *cj0700* (data not shown). This mutant phenotype was replicated in independent *cj0700* mutants constructed in other strains and could be complemented by the insertion of *cj0700* into a non-coding region between *cj0045c* and *mnmA*. Successful complementation *in trans* indicated that the mutant phenotype was due to the mutation of *cj0700* and not due to phase variation-based effects on motility (31) or polarity. Slight discrepancies in chemotaxis between wild type and complemented mutants may be expected as the genomic context of *cj0700* is different and the gene is being expressed constitutively from a *metK* promoter that has a low level of expression (32, 33). These differences may influence the expression of *cj0700*, however expression of *cj0700 in trans* from its own promoter gave the same pattern of complementation as the *metK* promoter (data not shown). Another confounding factor could be differing phase variation states between the wildtype and complementation mutant that could affect motility (31); the phasotypes of the strains were not determined (34). Nevertheless, the disruption of chemotaxis in a *cj0700* mutant mirrors similar studies in both *E. coli* and *H. pylori* where *cheZ* and *hp0170* mutants display impaired chemotaxis which can be complemented *in trans* (16, 24).

Having established a role for Cj0700 in chemotaxis in *C. jejuni*, protein-protein interaction with key Che proteins as well as the ability to dephosphorylate CheY-P was demonstrated. Cj0700 interacts specifically with Che proteins containing a response regulator domain. Pull down experiments showed that Cj0700 will bind CheY, CheA-RR and CheV. With the B2H assay the strongest interaction was between CheY and Cj0700, but both CheV and CheA-RR were also shown to interact with Cj0700, albeit at a lower level. There was no interaction with CheB, which in *C. jejuni* does not contain a RR domain. In *E. coli* CheB is not believed to be dephosphorylated by CheZ but loses phosphate through natural dephosphorylation (35).

In order to establish if Cj0700 can dephosphorylate CheY, an assay involving phosphorylation of CheY by CheA was developed. As CheY was found to rapidly auto-dephosphorylate under assay conditions at temperatures above 4°C, the assay was optimised to generate sufficiently stable levels of phosphorylated CheY. Purified Cj0700 was found to remove phosphate from CheY. Cj0700 dephosphorylation was not found to extend to other non-chemotaxis RR proteins as phosphorylated RacR (36, 37) is not dephosphorylated by Cj0700 (data not shown), suggesting the activity of Cj0700 is restricted to chemotaxis modulation. Therefore, we have experimentally verified that Cj0700 is a functioning CheZ homologue in *Campylobacter*. Apart from *H. pylori* (25) no other Cj0700/HP0170 homologue has been shown to have phosphatase activity on CheY. However, other CheZ-like phosphatases, such as the CheX and CheC/D found in *T. maritima* and *B. subtilis*, accelerate de-phosphorylation of CheY (38). Therefore, similar to *E. coli*, it is likely that *C. jejuni* chemotaxis depends on a combination of native dephosphorylation activity of CheY and the activity of a CheZ orthologue.

The *C. jejuni* CheA contains a RR domain, but it is not clear what role this domain has in chemotaxis. In *H. pylori*, deletion of the CheA-RR domain disrupts chemotaxis (39) and we have shown that the histidine kinase of CheA interacts with CheA-RR and phosphorylates it. At least *in vitro*, Cj0700 can remove the phosphate from CheA-RR, although not as effectively as from CheY. It is possible that this substantially lower level of de-phosphorylation is an artefact of the use of an isolated CheA-RR domain in an *in vitro* assay and is not relevant for intracellular CheA complexes.

CheV also contains a RR domain which CheA kinase can interact with and phosphorylate. When comparing RRs in the chemotaxis signaling pathway, the CheA-HK phospho-transfer rate to CheV was not as efficient as with CheY or CheA-RR domain. Similar effects have been seen in *H. pylori*, in which CheV phosphate uptake from CheA-HK was markedly reduced (40). Our phosphorylation experiments also indicated that Cj0700 could not efficiently remove phosphate from CheV under our assay conditions. The reduction in phosphate transfer from CheA-HK to CheV may have the same basis as the lack of affinity of Cj0700 with CheV -interaction between the W domain in CheV and the RR domain. Such an interaction may have a role in modulating coupling by CheV of chemoreceptors (41). CheV is likely to play an adaptive role in *Campylobacter* chemotaxis through interaction with the chemoreceptors (23). This function may be modulated through phosphorylation of the cognate CheV RR domain by CheA, but another protein, Cj0371, has also been shown to affect *C. jejuni* chemotaxis at least in part through interaction with CheV (42).

In conclusion, we have demonstrated, for the first time, that Cj0700, a remote orthologue of CheZ, is an important component of the signal transduction system that controls chemotactic motility of *C. jejuni*. We have also shown that CheA directly phosphorylates both the CheA-RR domain and CheV, albeit at a slower rate than the phosphorylation of CheY.

## Acknowledgements

The authors would like to thank Oliver Bridle and Esther Karunakaran for preliminary work that supported this study. Paul Ainsworth undertook the work to show CheA phosphorylation of CheY. Thanks to Duncan Gaskin (IFR, Norwich) and Arnoud van Vliet (University of Surrey) for plasmids used in the study, and Jo Wanford for help with drafting the manuscript. AJ would like to thank the Department of Genetics for support during his doctoral studies. BBSRC funding helped support this work (BB/J016667/1).

