## Supplemental material for "A CheZ orthologue in *Campylobacter jejuni* plays a role in chemotaxis through conserved phosphatase activity"

### Jama and Ketley Supplementary Material

#### Table 1S

##### *E. coli* strains and plasmids used in this study.

| Name | Description | Source/Reference |
|---|---|---|
| ***E. coli*** | | |
| XL1-Blue | *recA1, endA1, gyrA96, thi-1, hsdR17, supE44, relA1, lac,* [F´ *proAB, lacIqZΔM15* Tn*10* tet$^{R}$]. | Stratagene, UK |
| BL21 | F$^{-}$ *ompT hsdS*$_{B}$ ($r_{B}^{-}$, $m_{B}^{-}$) *gal dcm* (DE3) | New England Biolabs |
| Rosetta | Lactose permease (*lacY*) mutant, deficient in *lon ompT* proteases; contains plasmid encoding *argU, argW, glyT, IleX, leuW, metT, proL, thrT, thrU,* and *tyrU* | Novagen |
| BTH101 | Non-reverting adenylate cyclase deficient (cya) *E. coli* reporter strain for bacterial two-hybrid system. *F- cya-99 araD139 galE15 galK16 rpsL1 (Strr) hsdR2 mcrA1 mcrB1* | Euromedex, UK |
| **Plasmids** | | |
| pAJ4 | pUC19 containing 1896 bp insert with *cj0700* gene with 600 bp flanking sequence in plasmid. | This study |
| pAJ5 | pAJ4 containing 730 bp chloramphenicol resistance cassette from pAV110 cloned into remaining *cj0700* sequence. | This study |
| pAV110 | Cat cassette in pUC | (Amsterdam et al., 2003) |
| pAJ6 | Contains 896 bp *cj0700* fragment cloned into pKmetK46 | This study |
| pKmetK46 | Kanamycin resistance cassette (*aph3*) and divergent *metK* promoter both flanked by *cj0046* sequence. | Duncan Gaskin and (Reuter and van Vliet, 2013) |
| pLEICS03 | N-terminal polyhistidine tag vector with T7 promoter and TEV protease cleavage site (Km$^{R}$) | PROTEX, University of Leicester |
| pAJ8 | *cj0700* (696 bp) cloned into pLEICS03 | This study |
| pAJ7 | *cj0700*(D167N) (696 bp) in pLEICS03 | This study |
| pTrcHisB | N-terminal polyhistidine tag vector | Invitrogen |
| pPA016 | *cheV* (968 bp) cloned into pTrcHisB (Amp$^{R}$). | This study (Ainsworth) |
| pPA021 | *cheY* (430 bp) cloned into pTrcHisB (Amp$^{R}$). | This study (Ainsworth) |
| pPA017 | c*heA* (2323 bp) cloned into pTrcHisB (Amp$^{R}$). | This study (Ainsworth) |
| pPA024 | c*heA* histidine kinase domain (1906 bp) cloned into pTrcHisB (Amp$^{R}$). | This study (Ainsworth) |
| pPA025 | c*heA* response regulator domain (1972 bp) cloned into pTrcHisB (Amp$^{R}$). | This study (Ainsworth) |
| pPA037 | *cheY* (430 bp) cloned into pGex-4T-1 (Amp$^{R}$). | This study (Ainsworth) |
| pGEX-4T-1 | N-terminal glutathione tag vector | GE Healthcare |
| pEKG1 | *cj0700* (696 bp) cloned into pGex-4T-1 (Amp$^{R}$). | This study (Karunakaran) |
| pUT18 | Bait vector for two hybrid system (Amp$^{R}$). | Euromedex, UK |
| pKT25 | Prey vector for two hybrid system (Km$^{R}$). | Euromedex, UK |

| | | |
|---|---|---|
| pUT18:Zip | Two hybrid control plasmid containing leucine zipper region of the GCN4 yeast protein ($Amp^R$). | Euromedex, UK |
| pKT25:Zip | Two hybrid control plasmid containing leucine zipper region of the GCN4 yeast protein ($Km^R$). | Euromedex, UK |
| pKT25-CheY | *cheY* (417 bp) cloned into pKT25 vector ($Km^R$). | This study (Bridle) |
| pKT25-CheV | *cheV* (968 bp) cloned into pKT25 vector ($Km^R$). | This study (Bridle) |
| pKT25-CheB | *cheB* (575 bp) cloned into pKT25 vector ($Km^R$). | This study (Bridle) |
| pKT25-CheA-RR | c*heA* response regulator domain (1972 bp) cloned into pKT25 vector ($Km^R$). | This study (Bridle) |
| pUT18-Cj0700 | *cj0700* (722 bp) cloned into the pUT18 vector ($Amp^R$). | This study (Bridle) |

Cj0046* is here used as a label for a related region in NCTC11168, NCTC11828 and 81-176.

**Table 2. Primers used in this study**

| Primer | Sequence (5'-3') | Description |
|---|---|---|
| Cj0700F | CCCGGATCCGCGGCCGCCAGTATCAGCTATGCCACTATT | Amplification of cj0700 |
| Cj0700R | CCCGGATCCGCGGCCGCGCACAAGATCCACGATTGC | |
| Cj0700-invf-1 | CCCAGATCTCTTCTTGAGTCATTTTTTTCCCT | Inverse PCR mutagenesis |
| Cj0700-invr-2 | CCCAGATCTGCTTTAAGCCGTTATATGAGT | Inverse PCR mutagenesis |
| Cj0700D167N-F | ATGCAGTATCAAAATATACATCGTCAA | D167N |
| Cj0700D167N-R | TTGACGATGTATATTTTGATACTG | D167N |
| cj0700-F-pL03 | TACTTCCAATCCATGACTCAAGAAGAGCTTGATGCTTTG | pAJ8 |
| cj0700-R-pL03 | TATCCACCTTTACTGTCATCATTTTTGCCCTAAACTTG | pAJ8 |
| HCheY-F | GGAAGATCTGTGAAATTGTTAGTTGTTGATGACAGTTCTAC | BglII; pPA021 |
| HCheY-R | CGGGGTACCTTACTCAGCTGCACCTTCTCC | KpnI; pPA021 |
| HCheA-F | GGAAGATCTATGGAAGATATGCAAGAAATAC | BglII; pPA017 |
| HCheA-R | CGGGGTACCTCTTATCCTAGTTTCAAATTTTTTC | KpnI; pPA017 |
| HCheA-HK-F | GAAGATCTATGGAAGATATGCAAGAAATAC | BglII; pPA024 |
| HCheA-HK-R | GGTACCCCGTTAAATTGCATAGAATTCCTCTTGAG | KpnI; pPA024 |
| HCheA-RR-F | GGAAGATCTAGTTCATTTAAACTTAAAATTCCTCTTAC | BglII; pPA025 |
| HCheA-RR-R | GGTACCCCGTCATTACTCATATTCTTATCCTAGTTTC | KpnI; pPA025 |
| HCheV-F-XhoI | CCGCTCGAGAATGTTTGATGAAAATATCGTGAAAAC | pPA016 |
| HCheV-R-KpnI | CGGGGTACCTTACCCCTGTTCTTGAGATTGATG | pPA016 |
| Cj0700-GSTF | CCCGAATTCATGACTCAAGAAGAGCTTG | EcoRI; GST-tag |
| Cj0700-GSTR | CCCGTCGACTCATTTTTGCCCTAAACTTG | SalI; GST-tag |

**Supplementary Methods.**

**Bacterial strains, media and plasmids**

*C.jejuni* strains (Table 1) were routinely grown on Mueller-Hinton agar (MHA; Oxoid, Basingstoke, UK) and in Mueller-Hinton Broth (MHB Oxoid) supplemented with vancomycin (10 µg/ml) and trimethoprim (5 µg/ml) under microaerobic conditions (10% $O_2$ and 5 % $CO_2$ and 85% $N_2$) (Variable Atmosphere Incubator; Don Whitley Scientific Ltd, Shipley, UK) at 42 °C for between 2-5 days. Where necessary, mutants containing the respective antibiotic resistance cassette were grown on media supplemented with chloramphenicol (20 µg /mL), erythromycin (5 µg /mL) or kanamycin (50 µg /mL). *E. coli* (Table 1S) cells were grown on LA (Luria-

Bertani Agar), MacConkey agar plates or in liquid media (LB; Luria-Bertani Broth) supplemented with appropriate antibiotic selection.

**Chemotaxis and Growth Assays**

Basic cell motility of *C. jejuni* strains was determined by phase contrast microscopy. Strain chemotaxis was tested by spotting or stabbing 20 µl of an overnight cell suspension ($OD_{600}$ of 1.0) into 0.3% (w/v) semisolid MHA plates and incubated for 3-5 days at 42°C under microaerobic conditions.

*C. jejuni* growth patterns were determined either in individual tubes (inoculated at 0.05 $OD_{600}$ in 5 ml of media incubated at 42°C with shaking under microaerobic conditions), measuring $OD_{600}$ every 12 hours or in microtiter plates (200 µl of 0.05 $OD_{600}$ in each well with the plate sealed with gas permeable pre-pierced membrane and incubated overnight at 42°C with shaking under micro-aerobic conditions) where cell growth was measured every 3 hours post inoculation in a FLUOstar Omega microplate reader.

***C. jejuni* mutant construction.**

The NCTC11168 *cj0700* gene with flanking sequence was cloned utilising primers Cj0700F and Cj0700R (Table 2S) into the pUC19 *Bam*HI site (pAJ4; Table 1). The *cj0700* gene was mutated by insertion of a promoter-less and terminator-less chloramphenicol acetyltransferase (*cat*) gene from pAV110 (van Amsterdam et al., 2003) into a *Bgl*II site constructed by inverse PCR (Karlyshev et al., 1999) using primers Cj0700-invf-1 and Cj0700-invr-2 (Table 2S). For complementation of the *cj0700* mutation, *cj0700*, including the likely promoter region, was inserted utilising primers CCj0700_F and CCj0700_R (Table 2S), into the *Esp*3I site of pKmetK (Gaskin et al. 2007) (pAJ6 Table 1).

Mutated and complementing alleles were inserted into the genome of *C. jejuni* strains NCTC11168, NCTC11828 and 81-176 using natural transformation (Wang and Taylor, 1990) or electroporation (2.5 KV, 200 Ω and 25 µF). For mutation of cj0700, transformants in each strain were selected on MHA containing chloramphenicol to form AJ1 (NCTC11168), AJ4 (NCTC11828) and AJ6 (81-176) (Table 1). For complementation transformants were selected on MHA containing kanamycin to form AJ3, AJ7 and AJ10 respectively (Table 1), where the wild-type *cj0700* allele to complement the *cj0700*::*cat* mutation was inserted into the *cj0046* pseudogene between *cj0045c* and *mnmA* in NCTC11168 and the equivalent locus in other strains (Table 1). All mutations were verified by PCR and DNA sequencing. To control for loss of motility of NCTC11168, NCTC11828 and 81-176 following electroporation, the insertion mutation of *cj0700* and the insertion of *cj0700* into *cj0046* were transferred into motile variants of each strain by natural transformation and motility verified.

**Protein expression and purification**

His-tagged CheY, CheA, CheA-HK, CheA-RR and CheV were expressed using clones derived using pTrcHisB, and His-tagged Cj0700 using pLEICS03 (Table 1S). GST-tagged Cj0700 and CheY were expressed using pGEX-4T-1 (Table 1 and 2S). Tagged proteins were expressed in and purified from *E. coli*

Rosetta and BL21 strains. Log phase cultures were induced with 1 mM IPTG and incubated at 30 or 37 °C for 3 hours with shaking. GST tagged proteins were purified using GSTrap FF columns and His-tagged proteins using HisTrap FF columns (GE Healthcare Life sciences).

##### Phosphorylation/de-phosphorylation assays

Phosphorylation/de-phosphorylation assays were carried out using $\gamma^{32}$-P (3000 Ci/mmol and 10μCi/μl) ATP diluted with unlabelled ATP in reaction buffer (50 mM Tris-HCl (pH 7.5), 50 mM KCl, and 20 mM $MgCl_2$). CheA-HK was incubated with radio-labelled Pi for 10 minutes at 4, 30 or 42 °C prior to addition of RR domain-carrying proteins and Cj0700 proteins. At each time point, the reaction mixture was inactivated (2 x SDS sample loading buffer) and incubated on ice before separation by SDS-PAGE and dried. Visualization and quantification of protein interaction were made using a Typhoon 9400 Variable Mode Imager (Amersham Biosciences) and analysis was performed using the ImageQaunt TL software (GE Healthcare Life sciences).

##### Protein-protein interaction assays

GST-tagged protein (0.5 μg) was bound to glutathione Sepharose 4B resin beads, washed and His-tagged protein (0.5 μg) was added. Acetyl phosphate (30 mM) and 10 mM $MgCl_2$ were added (Appleby and Bourret, 1998, Zhao et al., 2002). Beads were re-suspended in 2x SDS sample buffer and proteins bound to the beads visualised by SDS-PAGE or Western blotting.

The bacterial two hybrid (B2H) assay for protein-protein interactions was undertaken using a system based on adenylate cyclase (AC) reconstitution (Battesti and Bouveret 2012) supplied by Euromedex. Bait (pUT18-based) and prey (pKT25-based) plasmids were transformed into *E. coli* BTH101 (Table 1S) and screened on MacConkey agar containing 1% (w/v) glucose-free maltose incubated at 30 °C for 3 days. Colour change was evaluated visually and interpreted as no interaction, weak interaction and strong interaction. The plasmids pUT18:Zip and pKT25:Zip were used as protein-protein interaction controls (Table 1S).


AMSTERDAM, K., VLIET, A. H. M., KUSTERS, J. G., FELLER, M., DANKERT, J. & ENDE, A. 2003. InducedHelicobacter pylorivacuolating cytotoxin VacA expression after initial colonisation of human gastric epithelial cells. *FEMS Immunology & Medical Microbiology,* 39**,** 251-256.

APPLEBY, J. L. & BOURRET, R. B. 1998. Proposed signal transduction role for conserved CheY residue Thr87, a member of the response regulator active-site quintet. *J Bacteriol,* 180**,** 3563-9.

KARLYSHEV, A., HENDERSON, J., KETLEY, J. & WREN, B. 1999. Procedure for the investigation of bacterial genomes: random shot-gun cloning, sample sequencing and mutagenesis of Campylobacter jejuni. *Biotechniques,* 26**,** 50-2, 54, 56.

REUTER, M. & VAN VLIET, A. H. 2013. Signal balancing by the CetABC and CetZ chemoreceptors controls energy taxis in Campylobacter jejuni. *PLoS One,* 8**,** e54390.

VAN AMSTERDAM, K., VAN VLIET, A. H., KUSTERS, J. G., FELLER, M., DANKERT, J. & VAN DER ENDE, A. 2003. Induced Helicobacter pylori vacuolating cytotoxin VacA expression after initial colonisation of human gastric epithelial cells. *FEMS Immunol Med Microbiol,* 39**,** 251-6.

WANG, Y. & TAYLOR, D. E. 1990. Natural transformation in *Campylobacter* species. *J. Bacteriol.,* 172**,** 949-955.

ZHAO, R., COLLINS, E. J., BOURRET, R. B. & SILVERSMITH, R. E. 2002. Structure and catalytic mechanism of the E. coli chemotaxis phosphatase CheZ. *Nat Struct Biol,* 9**,** 570-5.
